# Movement initiation and grasp representation in premotor and primary motor cortex mirror neurons

**DOI:** 10.1101/2019.12.17.880237

**Authors:** Jerjian S.J., Sahani M., Kraskov A.

**Author notes:** **Code Accessibility:** Codes and data to reproduce the analyses presented are available from the corresponding author upon reasonable request.

## Abstract

Pyramidal tract neurons (PTNs) within macaque rostral ventral premotor cortex (F5) and primary motor cortex (M1) provide direct input to spinal circuitry and are critical for skilled movement control, but surprisingly, can also be active during passive action observation. We recorded from single neurons, including identified PTNs in the hand and arm area of primary motor cortex (M1) (n=189), and in premotor area F5 (n=115) of two adult male macaques, while they executed, observed, or simply withheld (NoGo) reach-to-grasp and hold actions. We found that F5 maintains a more sustained, similar representation of grasping actions during both execution and observation. In contrast, although some M1 neurons mirrored during the grasp and hold, M1 population activity during observation contained signatures of a withholding state. This suggests that M1 and its output may dissociates signals required for the initiation of movement from those associated with the representation of grasp in order to flexibly guide behaviour.

**Significance Statement:** Ventral premotor cortex (area F5) maintains a similar representation of grasping actions during both execution and observation. Primary motor cortex and its outputs dissociate between movement and non-movement states.

## Introduction

The defining property of mirror neurons (MirNs) is that they modulate their firing both when a monkey performs an action, and when it observes a similar action performed by another individual (Gallese et al., 1996; Rizzolatti & Fogassi, 2014). Since their discovery in the macaque rostral ventral premotor cortex (F5), cells with mirror-like properties have been identified in parietal areas (Fogassi et al., 2005; Bonini et al., 2010; Lanzilotto et al., 2019), dorsal premotor cortex (PMd) (Cisek & Kalaska, 2004; Papadourakis & Raos, 2019), and even M1 (Tkach et al., 2007; Dushanova & Donoghue, 2010; Vigneswaran et al., 2013). MirNs thus appear to be embedded in an extended network (Bonini, 2016; Bruni et al., 2018), within a parieto-frontal circuit integral to the execution of visually-guided grasp (Jeannerod et al., 1995; Borra et al., 2017). The widespread activity within this circuitry during action observation takes place in the absence of detectable movement or muscle activity, despite the finding that even PTNs, which project directly to the spinal cord, can exhibit mirror properties (Kraskov et al., 2009; Vigneswaran et al., 2013).

F5 MirNs often show similar levels of activity during execution and observation (Gallese et al., 1996; Kraskov et al., 2009), however in M1-PTNs there is typically a reduced, or reversed, pattern of firing during observation relative to execution (Vigneswaran et al., 2013; Kraskov et al., 2014). By design, most action observation paradigms require movement suppression, and this disfacilitation of spinal outputs provides a rational, threshold-based explanation for why movement is not produced. However, there is substantial empirical evidence of both facilitation and suppression during movement execution in PTNs (Kraskov et al., 2009, 2014; Quallo et al., 2012; Vigneswaran et al., 2013; Soteropoulos, 2018) and putative pyramidal neurons (Kaufman et al., 2010, 2013), which suggests a more nuanced relationship between PTN activity and movement. At the spinal level, PTNs not only excite motoneurons via cortico-motoneuronal (CM) projections (Porter & Lemon, 1993; Rathelot & Strick, 2006), but also exert indirect effects via segmental interneuron pathways, which in turn display their own complex activity before and during movement (Prut & Fetz, 1999; Takei & Seki, 2013). A dynamical systems approach (Shenoy et al., 2013) has recently suggested that movement-related activity unfolds in largely orthogonal dimensions to activity during action preparation, such that movement is implicitly gated during movement preparation (Kaufman et al., 2014; Elsayed et al., 2016), and a similar mechanism has been hypothesised to operate during action observation (Mazurek et al., 2018). While the roles of F5 and M1 during the execution of visually-guided grasp have been studied extensively (Umiltá et al., 2007; Davare et al., 2008; Schaffelhofer & Scherberger, 2016), a more systematic understanding of the differences between action execution and observation activity in these two key nodes in the grasping circuitry could provide important insights into dissociations between representation of potential actions at the cortical level, and recruitment of descending pathways and muscles for actual action execution (Schieber, 2011). Along these lines, recent work comparing MirNs in premotor and motor cortex found premotor MirNs, but not those in M1, showed similar state transitions in execution and observation (Mazurek et al., 2018). State-space analyses have also previously found that F5 and the upstream anterior intraparietal area (AIP) exhibit different dynamics during immediate and delayed grasping actions (Michaels et al., 2018). Additionally, while previous work has examined grasp representation in F5 during inaction conditions (Bonini et al., 2014b), and reported little overlap between MirNs and neurons encoding self-action withholding, interleaved action and inaction within peri-personal space may provide a more ecologically valid framework for investigating movement suppression during action observation.

To explore the functional relationships between action execution, observation, and withholding, we compared the discharge of MirNs in M1 and F5 of two macaque monkeys, while they switched between executing, observing, and withholding reach-to-grasp and hold movements on a trial-by-trial basis. Electrical stimulation in the medullary pyramid was used to antidromically identify PTNs, and we leveraged the precise timing of task events within a naturalistic experimental paradigm to assess and compare the patterns of discharge of different populations of neurons across task conditions.

## Methods

### Monkeys

Experiments involved two adult male purpose-bred rhesus macaque monkeys (*Macaca mulatta*, M48 and M49, weighing 12.0kg and 10.5kg, respectively). All procedures were approved by the Animal Welfare and Ethical Review Body at the UCL Queen Square Institute of Neurology, and carried out in accordance with the UK Animals (Scientific Procedures) Act, under appropriate personal and project licences issued by the UK Home Office. The monkeys were single-housed based on veterinary advice, in a unit with other rhesus monkeys, with natural light and access to an exercise pen and forage area. Both monkeys gained weight regularly throughout the procedure. At the end of all experiments, both monkeys were deeply anaesthetised with an overdose of pentobarbital and perfused transcardially.

### Experimental task

In each session, the monkey sat opposite a human experimenter, with a custom-built experimental box apparatus between them (Figure 1A). The monkey was presented with two target objects in peri-personal space, a trapezoid affording precision grip (PG), and a sphere affording whole-hand grasp (WHG) (Figure 1A, inset). Each trial began after a short inter-trial interval (ITI) (1-2s), with the monkey depressing two homepads with both hands and the experimenter depressing a homepad on their side. A controllable LCD screen (14cm × 10cm) became transparent (LCDon, Figure 1B,C), and the object area was illuminated with white light. After a delay (0.25s in M48, variable 0.25-0.45s in M49), two amber LEDs illuminated on one side or the other to indicate the target object for the current trial. After a further delay (0.8s in M48, variable 0.8-1.2s in M49), a single green or red LED indicated the trial type. When a green LED was presented on the monkey side (Go), the monkey released the active (right) homepad (homepad release (HPR)), and made a reach-to-grasp movement towards the target object using their right hand. The monkey then grasped the object using a trained grasp (displacement onset (DO)), rotated the object into a window (> 30° rotation) and held for 1 second (hold onset (HO) to hold off). A constant frequency tone indicated that the monkey was in the hold window, and a second, higher frequency tone after 1s indicated successful completion of the hold. The monkey then released the object and returned to the homepad, and another high frequency tone indicated correct completion of the trial. Observation trials followed the same sequence, except that the experimenter performed the same reach-to-grasp and hold movement in front of the monkey, who remained still, with both hands on the homepads. On NoGo trials, a red LED required the monkey to simply remain on the homepads for the duration of the trial. After a delay (0.7s in M48, 1.0s in M49), a single tone indicated the end of the trial. The monkey received a small fruit reward directly to the mouth for each successfully completed execution, observation or NoGo trial. All trial types were presented in pseudo-randomised order, with relative proportions of 8:3:2 for each object. The larger proportion of execution trials were used to ensure the monkeys remained attentive and were regularly expected to move. Error trials, where there was a failure to respond appropriately within the constraints of the task (e.g. releasing the homepad before the Go cue), triggered a low frequency error tone and were immediately aborted by the experimental software. The monkey was not rewarded and these trials were excluded from further analysis.

**Figure 1.**
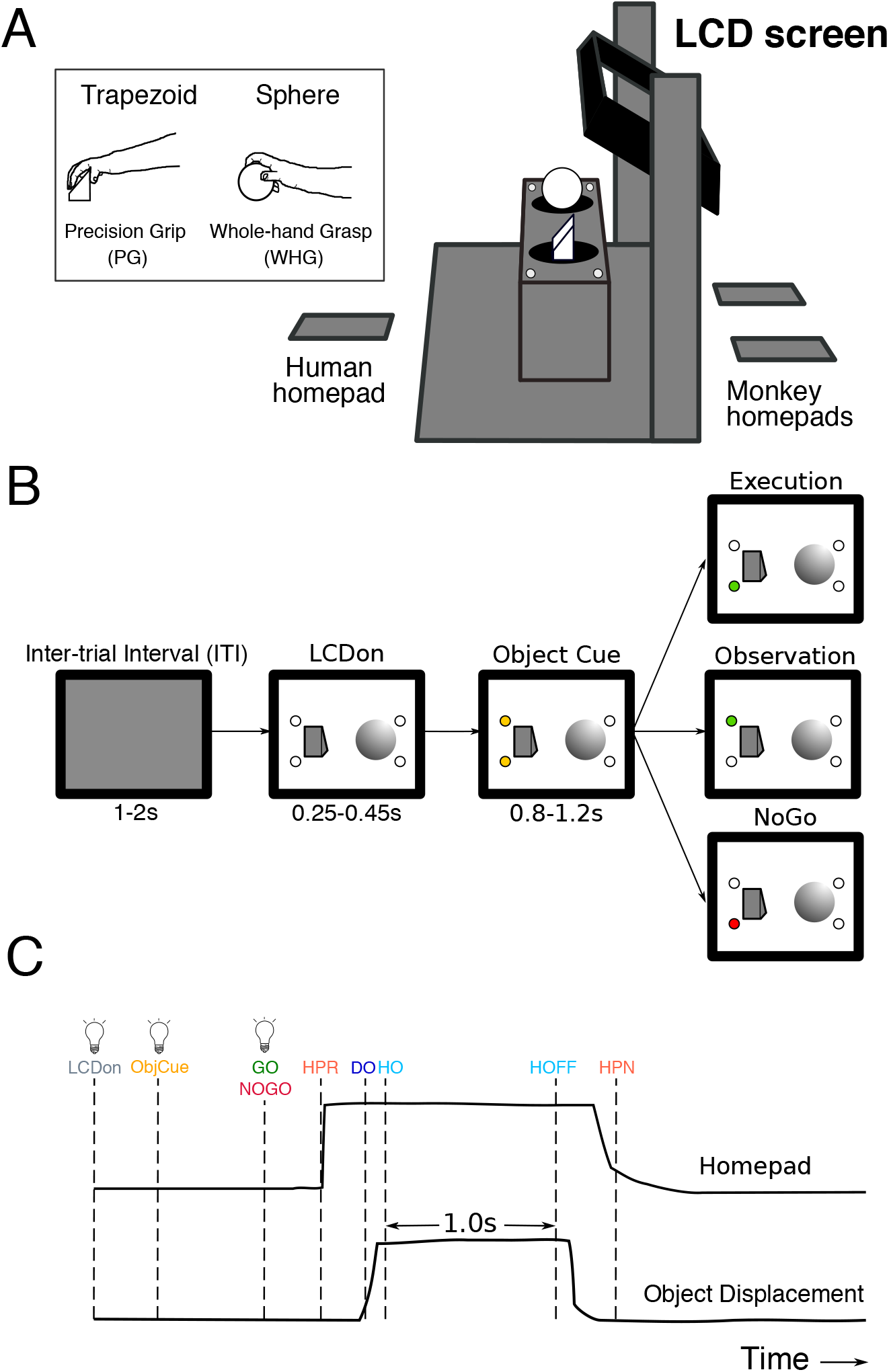
Basic mirror task design. **(A).** Schematic of the custom-built experimental box, showing target objects, their corresponding LEDs, LCD screen, and homepads. Inset shows the trapezoid and sphere objects, and the respective precision and whole-hand grasps performed by the monkeys on execution trials. **(B).** Pseudo-random trial presentation sequence, shown as 2-D representation of monkey’s view of object area. All trials began in the same way, with the object area illuminated (LCDon), and upcoming object/grasp cued (e.g. trapezoid, PG). Each trial was then indicated as Execution (green LED on monkey side), Observation (green LED on experimenter side), or NoGo (red LED on monkey side). **(C).** Homepad and object displacement signals on Go trials, and digital task events. **LCDon** LCD screen becomes transparent, **ObjCue**, object cue (amber LED); **Go/NoGo**, green/red LED; **HPR**, homepad release; **DO**, displacement onset; **HO**, hold onset; **HOFF**, hold offset; **HPN**, homepad return.

### Surgical implants

To prepare for recordings, subjects underwent several, well-spaced, surgical procedures under full general anaesthesia (induced with ketamine i/m 10mg/kg, maintained on 1.5-2.5% isoflurane in oxygen). First, a custom-designed TekaPEEK headpiece was secured to the skull for stable head fixation. In further surgeries, after the animal was fully trained, a) a TekaPEEK recording chamber was fixed with dental acrylic and bone cement to cover a craniotomy extending over primary and ventral premotor cortex; b) two tungsten stimulating electrodes were stereotaxically implanted in the left medullary pyramid c) subcutaneous recording electrodes were chronically implanted in up to 12 arm and hand muscles for electromyography (EMG) recording. After each procedure, animals were recovered overnight in a padded recovery cage, and received post-operative analgesic and antibiotics as prescribed under veterinary advice.

### Neuronal recordings

We used 16 and 7 channel Thomas Recording drives (Thomas Recording GmbH, Geissen, Germany), each containing 1–5 quartz glass-insulated platinum-iridium electrodes (shank diameter 80*μm*, impedance 1–2MΩ at 1kHz) to record in the arm/hand regions of M1 and F5. On a given recording day, we either carried out dual recordings, recording in M1 using the 16-drive, and in F5 using the 7-drive, or recordings in one area using a single drive. Linear array heads (spacing between adjacent guide tubes = 500*μm*) were used for initial mapping of M1 and F5, and subsequent recordings were conducted with square (16 drive) or circular (7 drive) heads to target more specific locations (305*μm* spacing). Penetration coordinates were estimated using a custom mapping procedure, based on triangulation of chamber lid coordinates measured in drive co-ordinates to an orthogonal system defined by stereotaxic coordinates of the same points measured during implantation of the recording chamber. Penetrations were made in the left (contralateral) hemisphere of each monkey, and aimed at the inferior bank of the arcuate sulcus (F5), and the hand/arm area of M1, just anterior to the central sulcus. Electrodes were independently lowered using custom computer software and adjusted in depth to isolate single unit activity as clearly as possible (Baker et al., 1999). Broadband signals from each drive were pre-amplified (x20, headstage amplifier), further amplified (x150), bandpass-filtered (1.5Hz – 10kHz), and sampled at 25kHz via a PCI-6071E, National Instruments card. We simultaneously recorded electromyographic activity from up to 12 muscles in the contralateral arm and hand, and analog signals of object displacement and homepad pressure (5kHz), as well as the precise timing of all task events at 25kHz resolution. All data was stored on laboratory computers for offline analysis. After recording at a site, repetitive intra-cortical microstimulation (rICMS) was delivered via an isolated stimulator. Sequences of 13 pulses at 333 Hz (duty cycle 0.5Hz) were delivered every 1-1.5s at intensities up to 30μA (M1), or 60μA (F5).

### PTN identification

While searching for cells, pyramidal tract (PT) stimulation was delivered between the two PT electrodes. The search stimulus intensity was 250–350*μ*A, and pulses were delivered every 0.6s (biphasic pulse, each phase 0.2ms). PTNs were identified as well-isolated cells which showed a robust and latency-invariant response (jitter ≤ 0.1ms) to PT stimulation. Double pulse search stimuli (separated by 10ms) were used to further help distinguish antidromic v.s. synaptic responses (Swadlow et al., 1978). We recorded the antidromic latency of each PTN, determined threshold, and used discriminated spontaneous spikes to collide the antidromic response, providing unequivocal identification of a PTN. PTN identification was always performed before task recordings, so this sample of cells was unbiased in terms of task-related activity.

### Spike discrimination

Offline spike sorting was performed using modified WaveClus software (Quiroga et al., 2004; Kraskov et al., 2009). Broadband data was first high-pass filtered (acausal 4th order elliptic 300Hz-3kHz, or subtraction of a median-filtered version of the signal). Threshold crossings were then sorted into clusters using an extended set of features, including wavelet coefficients, amplitude features, and the first 3 principal components. PTN spike shapes during task recordings were compared to the recorded waveforms of spontaneous spikes which resulted in successful collisions (Lemon, 1984; Kraskov et al., 2009). Single units were considered as those with a clean, consistent waveform and with inter-spike interval histograms uncontaminated below 1ms for bursting units.

## Data analysis

### EMG and behavioural analysis

EMG data for each channel was high-pass filtered (30Hz, 2nd order Butterworth), rectified, low-pass filtered (500Hz, 2nd order Butterworth), downsampled to 500Hz, and smoothed with a 100ms moving average. Signals were then aligned to the Go cue on individual trials, normalised to the 99th percentile amplitude across all trials and then averaged across trials within each condition. We recorded the timing of all relevant task events for subsequent alignment to analog signals. We defined reaction time on each execution and observation trial as the time between the GO cue and HPR, and movement time as the time between HPR and DO. For visualisation of displacement and homepad signals (Figure 2), individual trials were aligned to the Go cue. Signals were normalised to the 99th percentile amplitude across all trials and then averaged across trials within each condition.

**Figure 2.**
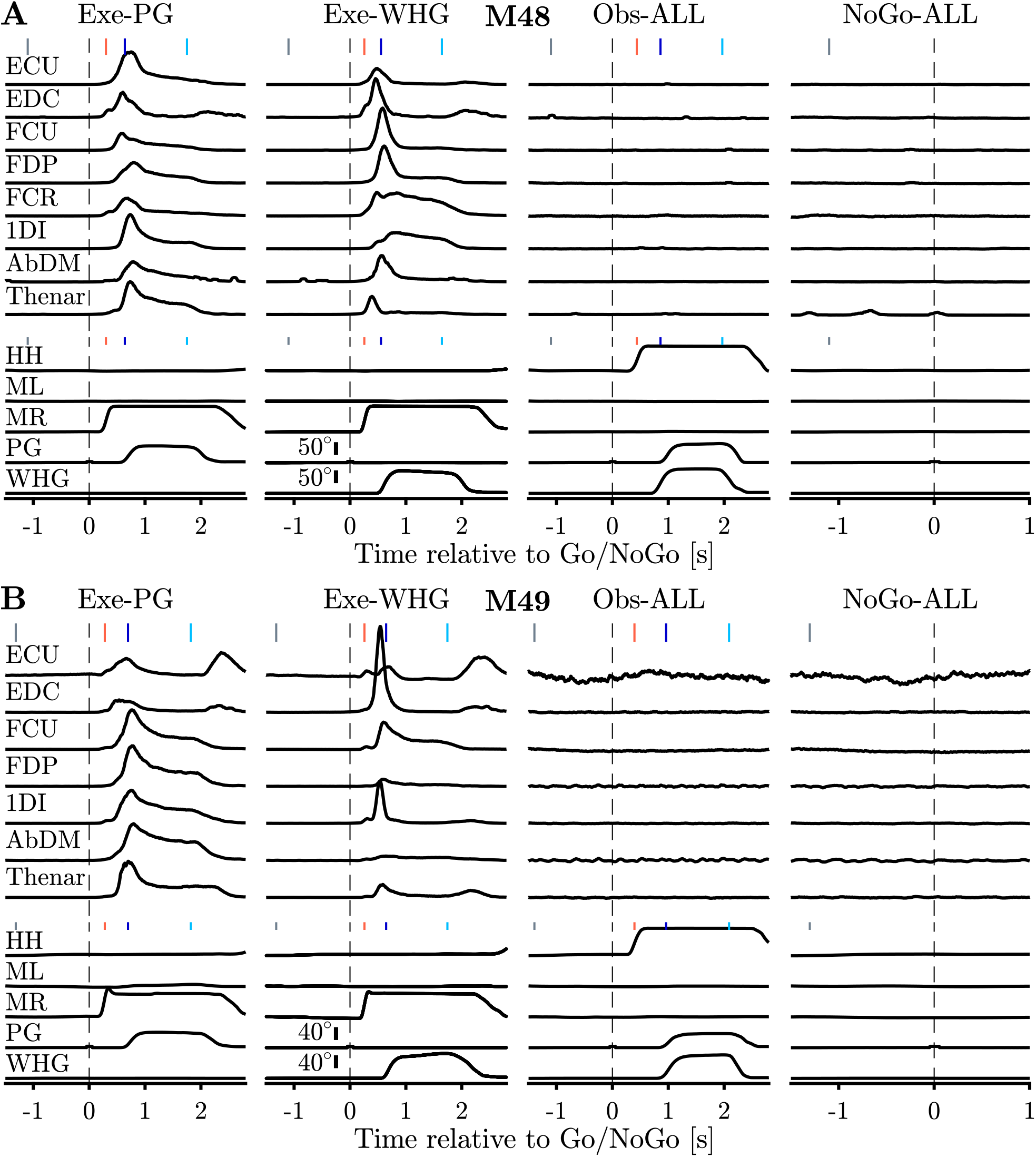
EMG and behaviour during basic mirror task. **(A).** A single session in M48. Top panels show pre-processed, rectified, and normalised EMG activity for different muscles with clean recordings. Corresponding object displacement and homepad signals are shown in bottom panels. Execution EMG is presented for both objects separately, observation and NoGo EMG are pooled across objects, and shown with a 10x higher gain. Vertical markers at top of each trace indicate median time of key task events (LCDon, HPR, DO, HOFF) relative to Go/NoGo cue (vertical dashed lines). **(B).** Same as (A), but for M49. **ECU**, extensor carpi ulnaris; **EDC**, extensor digitorum communis; **FCU**, flexor carpi ulnaris; **FDP**, flexor digitorum profundus; **FCR**, flexor carpi radialis; **1DI**, first dorsal interosseous; **AbDM**, abductor digiti minimi; **HH**, human homepad; **ML**, monkey left homepad; **MR**, monkey right homepad; **PG**precision grip; **WHG**, whole-hand grasp.

### Single-neuron analyses

To assess task-dependent modulation during execution and observation, we initially defined seven task epochs of interest, as follows. (1) LCDon-CUEon (2) Object Cue: 500ms period before the Go/NoGo cue. (3) Early React: 0-150ms from the Go/NoGo cue (4) Late React: 150-300ms from the Go/NoGo Cue. (5&6). Early and Late Reach: the first and second halves of the HPR-DO interval, which varied in length on each trial. (7) Hold: 0-700ms from HO. Firing rates during execution and observation for each neuron and object were subjected to a one-way ANOVA with factor EPOCH (7 levels), followed by post-hoc comparisons to Baseline (Tukey-Kramer correction for multiple comparisons). Neurons showing a significant ANOVA result and at least 1 significant post-hoc result (p < 0.05) for a given condition, with a raw firing rate range of 5 spikes s^−1^ in at least one condition, were considered to be task-modulated. Neurons modulated during any of epochs 5-7 (Movement epochs) of execution and observation were classified as MirNs. All modulation statistics and analyses were performed within object – a neuron could therefore be classed differently for the two objects. For some analyses, we further categorised MirNs according to the sign of their maximum modulation during the Movement epochs of both execution and observation, for each object separately. Thus, MirNs could be subdivided into facilitation-facilitation (F-F), facilitation-suppression (F-S), suppression-suppression (S-S), or suppression-facilitation (S-F) types for each object, based on their responses to execution and observation, respectively.

### Population analyses

For all population analyses, spike times for each neuron were binned into firing rates, baseline-corrected and normalised. The exact details differed for different analyses, and these are described in turn below.

#### Heatmaps and population averages

To visualise neural population activity during the task, spike counts in 10ms bins were smoothed with a Gaussian kernel (unit area, standard deviation 50ms) and converted to spikes s^−1^. As the timing of events varied across trials, conditions and sessions, firing rates were aligned separately to Go, HPR, and DO events on each execution and observation trial as appropriate, so that the relative timing of these three events, covering the most dynamic period of the task, was matched across all conditions and units. For visualisation purposes, peri-stimulus time histograms (PSTHs) aligned to different task events were interpolated to produce one continuous firing rate for each condition. The Go/NoGo event was set as time 0, and HPR and DO were defined as the mean times across conditions, objects, and sessions. The average firing rate across conditions during the Baseline period (LCDon-ObjCue) was subtracted, and the resultant net firing rates were soft-normalised to the maximum absolute firing rate across all conditions + constant factor of 5. Each unit’s firing rate across all conditions was therefore limited to a maximum theoretical range of [−1,1], where negative normalised values correspond to suppression of the firing rate relative to the baseline (Kraskov et al., 2009; Vigneswaran et al., 2013). Comparisons of population activity were conducted on firing rates within the intervals defined for single-neuron analyses (also baseline-corrected and soft-normalised), using Wilcoxon sign-rank tests with Bonferroni correction for multiple comparisons.

#### Correlation analyses

To make an initial analysis of the correspondence between execution and observation activity across the task, we averaged each neuron’s activity within each of the 7 task periods, and then across trials, for each condition. Activity was baseline-corrected by subtracting the average activity in the 250ms prior to LCDon, and then soft-normalised by the maximum absolute rate across conditions (within object), with a small constant (+5) added to the denominator to reduce the influence of low-firing neurons and improve interpretability of scatter plots. For each epoch, the net normalised execution and observation activity within a MirN population were extracted as N-dimensional vectors (N = number of MirNs), and the Pearson correlation coefficient between pairs of vectors was calculated. To compare observed correlation values to those expected by chance, we repeatedly shuffled (1000 iterations) the observation vector to destroy any within-unit relationships, and re-calculated the correlation coefficient, generating a null distribution of correlation values. Observed correlations were deemed significant if they fell beyond the range of 95% of the values in the null distribution. To examine the stability of cross-condition similarity in each population, we extended the cross-condition correlation procedure to correlate activity across timepoints, using time-resolved firing rates. To avoid trivial correlations induced by Gaussian smoothed firing rates, we calculated spike rates in 50ms non-overlapping bins, with the same multiple alignment as used for the population averages (Go, HPR, DO). We then correlated PSTH activity at execution condition timepoint *t* with activity at all timepoints *t* = 1…*T* in the observation condition, and vice versa, and then averaged across the diagonal. This produced a *T × T* matrix containing the correlation values of each timepoint *t* with every other timepoint.

#### Decoding analyses

We used the Neural Decoding Toolbox (Meyers, 2013) to examine how well activity in each sub-population discriminated between conditions. We first ran the decoding across all three conditions (Execution, Observation, NoGo), and then repeated analysis using Observation and NoGo conditions only. Binned data (non-overlapping 50ms bins), singly aligned to the Go/NoGo cue for each trial, was used to form pseudo-populations of units for each population separately, using 10 trials from each condition (3×10 = 30 data points for each condition in the 3-way decoding), and then randomly grouped into 10 cross-validation splits (3 data points per split). Firing rates were z-scored to reduce the bias of high-firing units in the classification. A maximum correlation coefficient classifier was trained on all but one of the splits, and then tested on the left-out split, and this procedure was repeated up to the number of splits, leaving out a different split each time. For increased robustness, the cross-validation splits were resampled 50 times, and decoding accuracy was averaged across these runs. To assess the significance of the observed decoding accuracy, we used a permutation test procedure. The classification was performed exactly as for the original data, except the relevant trial condition labels were shuffled beforehand. This was repeated 50 times to generate a null distribution of the decoding expected by chance, and the observed decoding accuracy was considered significant for a given bin if it exceeded all the values in the null distribution. To reduce the false positive rate, bins were considered truly significant only if they fell within a cluster of at least 5 consecutive significant bins.

#### Subspace analyses

To compare the trajectories of MirN activity in each sub-population, we applied principal component analysis (PCA). PCA identifies an orthogonal transformation for (correlated) data, where each successive dimension in the transformed space captures the maximum possible variance in the data, while remaining orthogonal to all other dimensions. Projection of data onto the leading principal axes can therefore be used to reduce dimensionality in a principled manner, and reveal low-dimensional structure which may otherwise be obscured. To apply this method to our data, PSTHs (firing rates in 10ms bins, convolved with a Gaussian kernel of unit area and 50ms standard deviation) were used to form pseudo-population firing rate matrices for each condition and neuronal sub-population. To prevent high-firing neurons from dominating the analysis, but preserve some relative range of firing rates, firing rates were soft-normalised by the total firing rate range across all times and conditions (+ a small constant of 5 spikes s^−1^).

Trial-averaged execution data from 50ms before the HPR cue to 500ms after HO, separately for each object, was then used to form a peri-movement activity matrix **M** (*T × N*, where *T* was the number of timepoints and *N* was the number of MirNs), which was then centred by subtracting the mean activity across time for each neuron (dimension). We projected trial-averaged execution and observation data spanning this time period onto the first *k* principal axes (*k* = 3; 3 dimensions typically captured >90% of the variance in **M**), yielding k principal components for each condition, each with a fractional variance associated with it. We quantified the overlap, or “alignment” of observation activity within this space by normalising the total captured variance by the maximum observation variance which could be captured by *k* axes, according to the following equation (c.f. Elsayed et al., 2016).

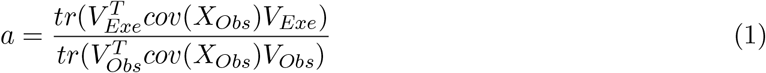

**V**_**Exe**_ and **V**_**Obs**_ are the first *k* eigenvectors of **X**_**Exe**_ and **X**_**Obs**_, where **X**_**Exe**_ and **X**_**Obs**_ are the mean-centred execution and observation activity, respectively. *tr* denotes trace. The denominator is mathematically equivalent to the sum of the eigenvalues of the first *k* eigenvectors of **X**_**Obs**_ and the alignment index is thus bounded between 0 (if **X**_**Exe**_ and **X**_**Obs**_ are fully orthogonal) and 1 (if **X**_**Exe**_and **X**_**Obs**_ are perfectly overlapping). We compared true alignment values to a null distribution of alignment of 10,000 pairs of random, orthonormal subspaces, and a p-value was computed as the proportion of values in the null distribution greater than the true alignment. P < 0.05 was considered significant (i.e. the true alignment value exceeded 95% of the values within the null distribution).

To examine the state-space overlap between observation and NoGo, we used PCA to define a second set of 3 principal axes using trial-averaged observation data from across all neurons, 100-400ms after the Go cue. We then projected activity from all three conditions onto these axes, and quantified variance captured and alignment statistics in an analogous way to that for the movement period subspaces.

## Results

We recorded single neurons in F5 and M1 of rhesus macaques performing and observing reach-to-grasp and hold actions, and assessed the profiles of activity across conditions and populations. We then compared modulation during the action observation condition, where monkeys were required to remain still, to neural activity when monkeys were explicitly cued to withhold movement.

### EMG and behaviour

Monkeys were trained to a high level of performance before recording (>90% correct trials per session). For both monkeys, reaction and movement times were significantly faster than human experimenters (Table 1, Wilcoxon sign-rank test on session averages, all p < 1 × 10^−13^). As the trapezoid object was positioned contralateral to the reaching (right) arm, monkey movement times were 30-50ms longer than those for the sphere (Wilcoxon sign-rank test, both monkeys p < 1 × 10^−7^). To verify that neural activity during action observation and withholding was not confounded by muscle activity, we simultaneously recorded EMG from up to 12 hand and arm muscles. During action execution, we observed characteristic patterns of EMG for each grasp. In the action observation and NoGo conditions, on the other hand, EMG activity was negligible (Figure 2, observation and NoGo are plotted at ×10 gain).

**Table 1.** Behaviour during recording sessions for basic mirror task. **RT**, reaction time; **MT**, movement time. Mean±SEM of median values from each session, rounded to nearest millisecond.

|  | M48 |  |  |  | M49 |  |  |  |
| --- | --- | --- | --- | --- | --- | --- | --- | --- |
|  | Monkey |  | Monkey |  | Monkey |  | Monkey |  |
|  | PG | WHG | PG | WHG | PG | WHG | PG | WHG |
| RT (ms) | 310±25 | 267±22 | 469±38 | 442±44 | 272±22 | 268±16 | 412±48 | 401±41 |
| MT (ms) | 306±20 | 279±14 | 430±31 | 374±38 | 404±23 | 351±20 | 520±39 | 532±45 |

### Effects of repetitive intracortical microstimulation

We delivered rICMS at 57 sites containing M1-PTNs, 124 sites with unidentified neurons (UIDs) in M1, and 111 sites in F5. Finger or thumb effects were elicited at 27/57 M1-PTN sites, 89/114 M1-UID sites, and 75/111 F5 sites. The majority of these sites in M1 had low thresholds (20/27 (74.1%) and 76/89 (85.4%) ≤20*μA*, PTNs and UIDs respectively), but not in F5 (27/75 (36.0%)).

### Database

Single neurons were recorded across 25 sessions in M48, and 40 sessions in M49. Across the two monkeys, we recorded a total of 304 neurons for ≥10 trials per object for both execution and observation conditions (Table 2), and 302 of these were also recorded for ≥7 NoGo trials per object. 189 units were recorded in M1, and 115 in F5. 60 M1 neurons were identified as PTNs; the remaining 129 were UIDs. F5-PTNs were observed and recorded, however total numbers were low (15 in M48, 8 in M49), so all F5 neurons were considered as one population. Figure 3 shows an MRI rendering of example penetrations in M48 in M1 and F5.

**Table 2.** Number of single-units recorded in each monkey and sub-population for at least 10 execution and 10 observation trials per grasp.

|  | M48 | M49 | Total |
| --- | --- | --- | --- |
| M1-PTN | 36 | 24 | 60 |
| M1-UID | 78 | 51 | 129 |
| F5 | 72 | 43 | 115 |
| Total | 186 | 118 | 304 |

**Figure 3.**
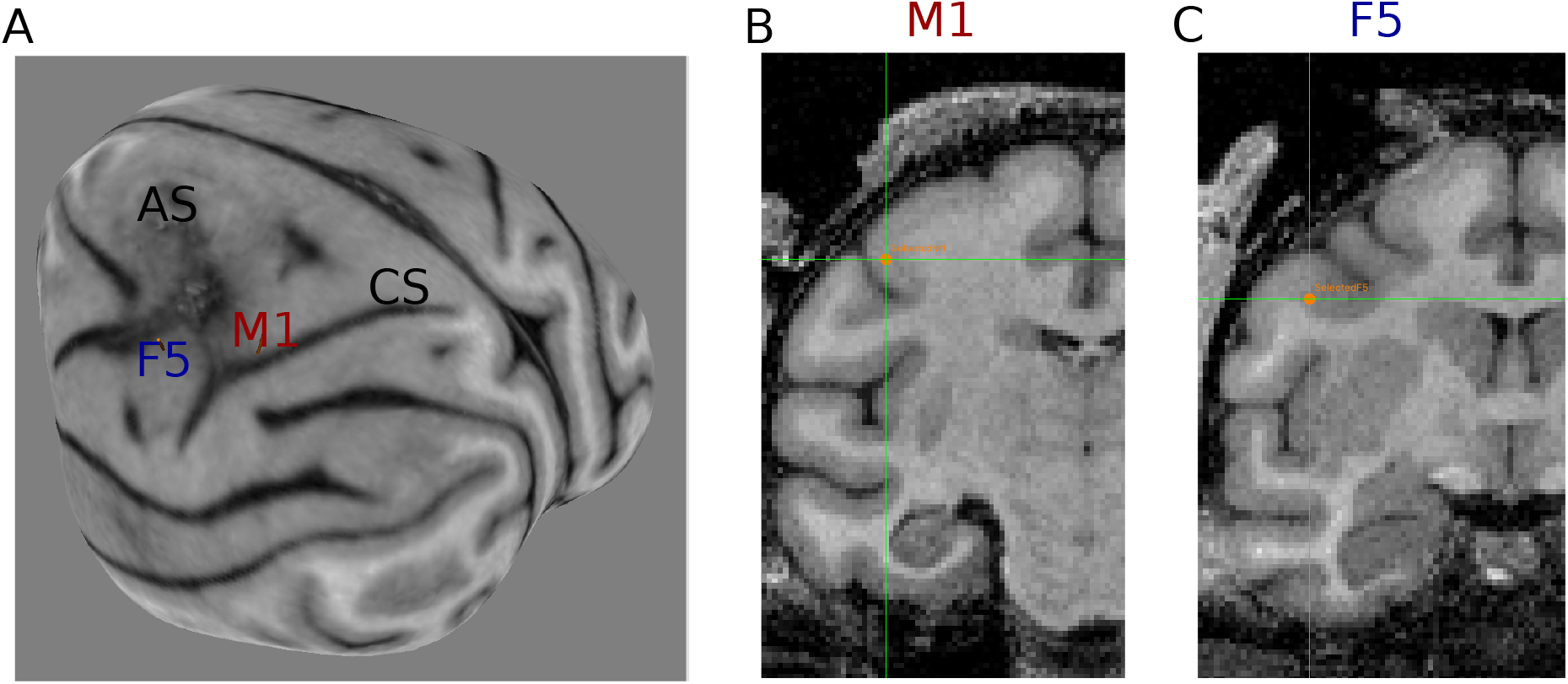
Structural MRI and example penetrations in M48. **(A).** shows a 3-D rendering of the brain surface. **CS**, central sulcus; **AS**, arcuate sulcus. **(B). & (C).** show coronal sections marking the same penetrations shown in (A) in M1 and F5, respectively. An additional transformation was required to translate between stereotaxic and MRI co-ordinates.

### Single-neuron responses

The complex naturalistic task set-up evoked a wide variety of responses in recorded neurons, particularly during action execution. A substantial proportion of neurons also showed responses to one or both of the action observation and NoGo conditions. Figure 4A shows a PTN recorded in M1, which maintained a steady baseline firing rate until the Go cue. During execution, HPR was accompanied by a suppression of firing for both grasps, followed by an increase leading up to DO, which was greater for WHG. This increased firing was maintained during the hold period, before falling below baseline as the monkey returned to the homepad. During observation, firing rates were considerably lower than during execution, and no suppression was apparent at movement onset. Firing rates increased to almost twice the baseline level during the hold period, before gradually decreasing, and were more similar across grasps than in execution. Another M1-PTN (Figure 4B) had a complex pattern of activity during execution, with a peak just before DO, particularly for PG, and a second peak prior to release of the object. During observation, the same unit showed a small, sustained increase in firing rate. The unit shown in Figure 4C was recorded in F5. In both action execution and observation conditions, the HPR to DO period showed a dramatic increase in firing for both grasps, peaking at >100 spikes s^−1^. In execution, hold period activity then stabilised at a lower rate, with activity for WHG sustained at a higher level. By contrast, observation activity decayed back to baseline relatively quickly at the beginning of the hold period. Figure 4D shows an M1-PTN with a steady baseline firing rate, which completely silenced during both execution and observation hold period, before showing some rebound at the end of the hold period, particularly during execution.

**Figure 4.**
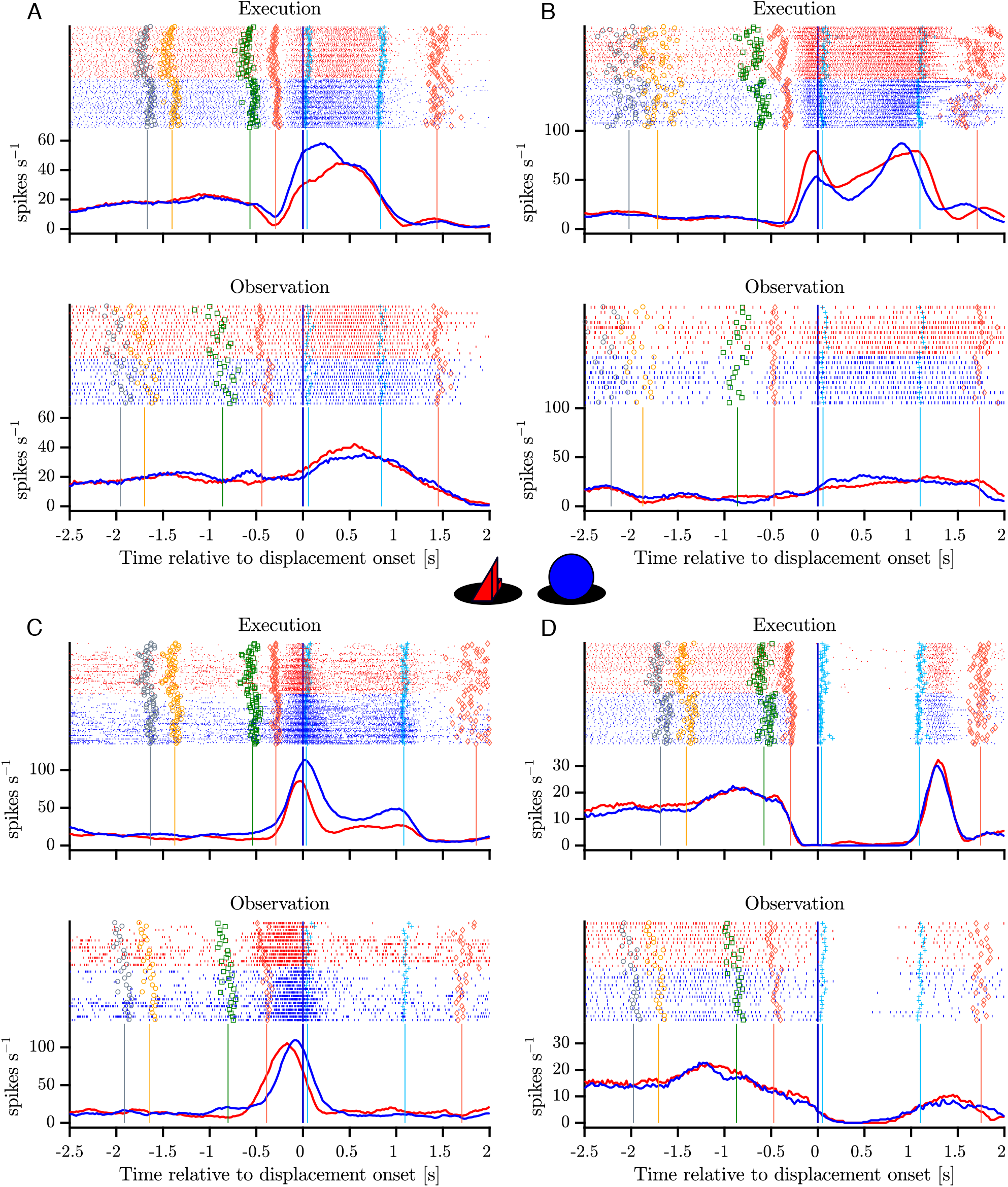
Example mirror neurons in M1 and F5. Raster and histogram representations of single neuron activity during execution (top panels) and observation (bottom panels). Activity is aligned to object displacement (DO). Rasters are split by grasp (PG and WHG, objects shown in inset) and condition for easier visualisation, although trials were presented in a pseudo-randomised order during recording. **(A-C).** Facilitation responses during execution and observation in two M1-PTNs and F5 unit, respectively. **(D).** M1-PTN showing pronounced suppression of activity during both execution and observation. Units in (A), (C) and (D) were recorded in M48, (B) was recorded in M49. Single trial events are indicated on raster plots (LCDon, Object Cue, Go, HPR, HO, HOFF, HPN), and median times relative to alignment are shown on histograms. Event colours are as shown previously (Figure 1C): LCDon – grey; Object Cue – orange; Go – green; HPR & HPN – magenta; HO & HOFF – cyan). For histograms, firing rates were calculated in 20ms bins and boxcar-smoothed (200ms moving average).

### Population-level activity during execution and observation

Although the connections of most of the recorded neurons were unknown, it is clear that action execution and observation produced different and complex response patterns both within and across units. For each neuron and task condition, we first assessed the statistical significance of changes in firing rate across relevant task intervals. For PG, 222/304 neurons (73.0%) showed significant modulation relative to baseline during the movement periods of execution (1-way ANOVA and post-hoc comparisons; see Methods). 117/222 (52.7%, 38.5% of total) were also modulated during action observation movement periods, and were therefore classed as MirNs. For WHG, 235/304 (77.3%) modulated during execution movement, and 110 of these (46.7%, 36.2% of total) were MirNs. Grasp specificity was frequently apparent during execution, whereas observation responses, when present, were often more comparable across the two objects (Figure 4), consistent with the broad congruence found in many MirNs (Gallese et al., 1996).

The extent of modulation during action observation may differ across premotor and motor cortex at the population level, with important implications for the effect of this activity on downstream targets. The heatmaps in Figure 5A-C show the time-resolved net normalised firing rate during execution and observation across the three MirN sub-populations, and Figure 5D-F show the averages during execution and observation for the facilitation-facilitation and facilitation suppression units. Within each sub-population, we found both facilitation and suppression responses relative to baseline during execution and observation, and the relationship between activity in the two conditions was variable. For the commonest group of identified MirNs, net normalised activity of facilitation-facilitation MirNs (those which increased their activity during execution and observation) was generally larger during execution movement than observation (Figure 5D-F, top panels). Population activity during the hold period of execution was substantially larger than that during observation in M1-PTNs (Wilcoxon sign-rank tests with Bonferroni correction for multiple comparisons, p = 0.0011 for PG, p = 0.011 for WHG), but not in M1-UIDs or F5 (all p > 0.05). To examine potential differences in time-varying pattern of activity during action execution and observation, we computed the correlation between execution and observation activity across all MirNs during different task epochs (Figure 6). During ObjCue, when trials were identical from the monkey’s perspective, all populations as expected showed a strong, significant correlation (r > 0.8, p < 1 × 10^−13^) between the two conditions (Figure 6A, top row). Contrastingly, activity patterns during the early stages of the reach were markedly different (Figure 6A, middle row). This was particularly the case in M1-PTNs, which showed no significant relationship between execution and observation activity at this stage of the task (r = 0.23, p = 0.2). M1-UIDs and F5 populations were also less well correlated during this period than before the Go cue, although the correlations remained significant (p < 0.0005). During the Hold period, activity patterns across the conditions became significantly correlated again during the grasp/hold stages of the task (p < 0.0001, Figure 6A, bottom row). We also compared the observed correlation values to a null distribution created by shuffling the observation vector so that within-unit relationships were lost. Correlations during the early reach period were significantly greater than all values in the null distribution for M1-UIDs and F5 (both grasps, p = 0), but not M1-PTNs (both grasps, p > 0.1, Figure 6B). For WHG, M1-PTN execution and observation activity during Late React and Late Reach epochs were also uncorrelated (p > 0.05). To assess the temporal stability of cross-condition similarity, we performed a cross-temporal pattern analysis using time-resolved PSTHs, by computing the correlation between net normalised activity at each timepoint with that of every other timepoint (Figure 7). The diagonal of this matrix therefore roughly corresponds to the epoch-based correlation values above. Activity prior to the Go cue, and during the hold period, was generally well correlated across the two conditions in all three populations and for both grasps. F5 neurons showed stronger correlations between the object cue and later hold periods, which was not apparent for M1-PTNs, indicating that the pattern of activity in these two periods was more consistent in F5.

**Figure 5.**
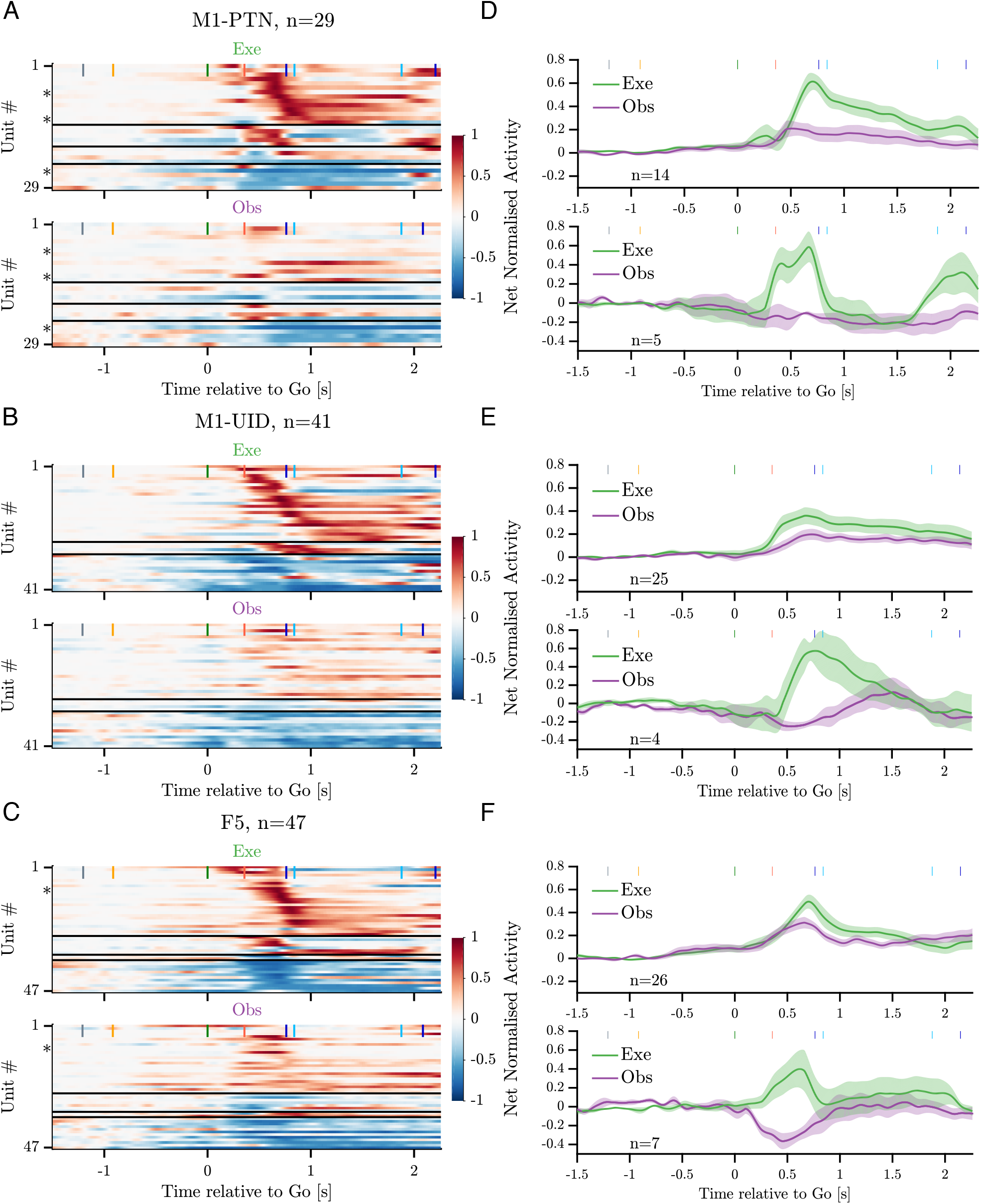
Mirror neuron population activity. **(A-C).** Heatmaps of net normalised activity in MirNs of each sub-population. Neurons are split into facilitation-facilitation, facilitation-suppression, suppression-facilitation, and suppression-suppression categories based on the sign of their modulation during action execution and observation relative to baseline. Horizontal black lines indicate splits between categories. Within each category, neurons are sorted based on the latency of their absolute peak response during execution (peak calculated between GO and HO+0.5s). Asterisks denote units shown in Figure 4. **(D-F).** Population averages for F-F (top panel) and F-S categories (bottom panel) separately for each sub-population.

**Figure 6.**
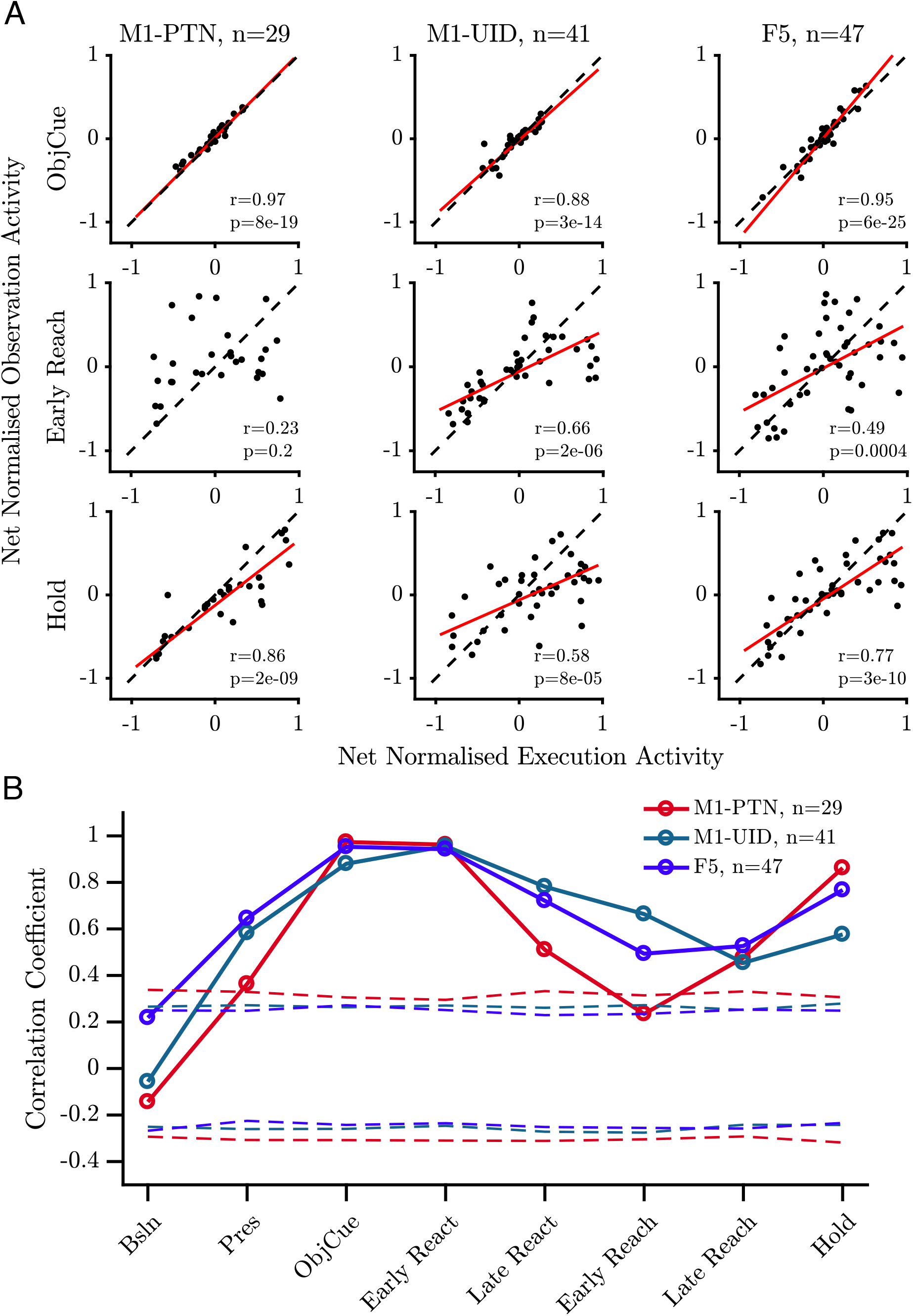
Relationship between execution and observation activity. **(A).** Net normalised activity during PG observation plotted against execution for the ObjCue, Early Reach, and Hold epoch, for MNs in each population (M1-PTNs left, M1-UIDs middle, F5 right). Pearson correlation coefficient r and corresponding p-value are shown in the lower right of each panel. Line of best fit for significant correlations are shown in red, dashed black traces mark *y* = *x* line. **(B).** Summary of Pearson correlation coefficients between execution and observation for each MirN population and PG task epoch. Dashed coloured lines denote 5 and 95th percentile correlation coefficients derived from 1000 shuffles for each population.

**Figure 7.**
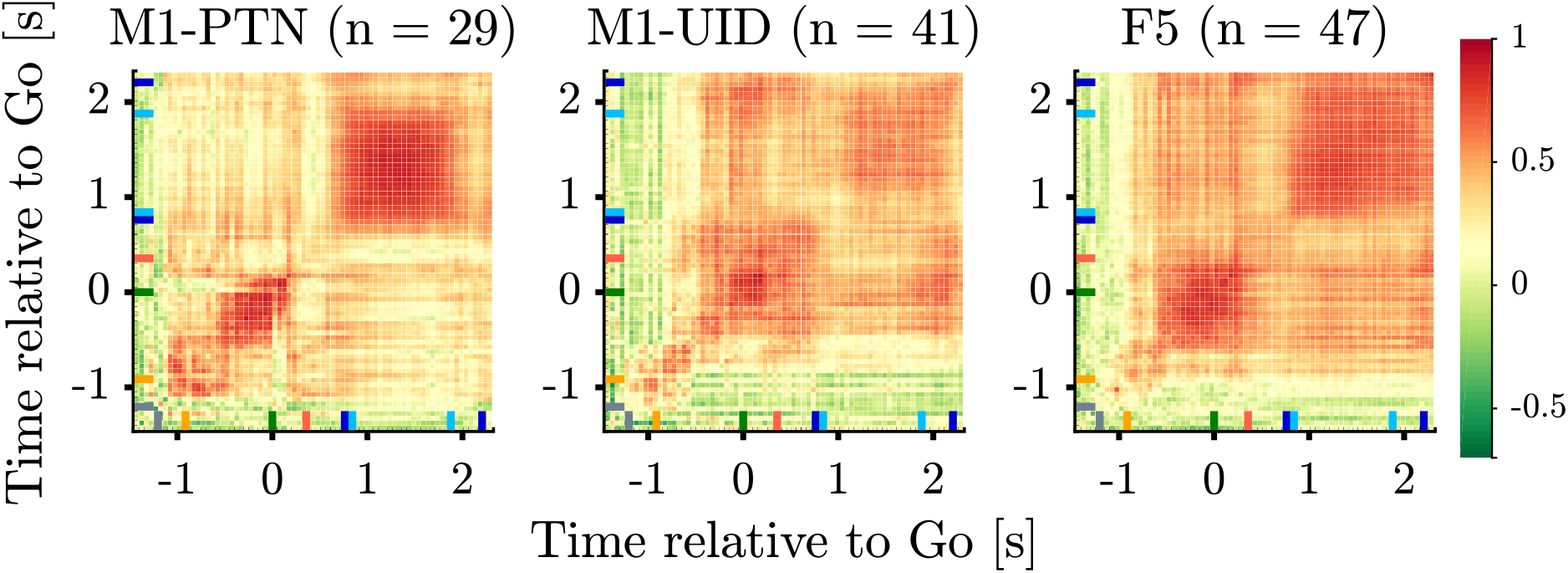
Cross-temporal correlation between execution and observation. Correlation across time between MirNs in each sub-population during execution and observation for PG (top), and WHG (bottom), M1-PTNs (left), M1-UIDs (middle), F5 (right). Green colours indicate low correlations, darker red shades indicate high correlation. Coloured ticks along left and bottom of each panel denote average event times across conditions, colour codes as in Figure 4 & Figure 1C.

We next used PCA to examine the nature of time-varying patterns of activity across action execution and observation in each sub-population within a movement subspace. PCA identifies a few dominant modes, or dimensions of neural activity within the full dimensional space which capture the majority of the variance in the data. The activity of the same neurons recorded during a different behaviour or time period can then be compared to the first based on the similarity of the covariance across neurons, which will result in similar or different projections upon the defined dimensions. This holds advantages over unweighted averaging of neural activity in different conditions, which also reduces dimensionality, but altogether sacrifices information regarding the relationships between different neurons and conditions. We defined a movement subspace empirically for each sub-population, using trial-averaged activity during execution reach and grasp, and then visualised evolution of execution (green) and observation (purple) trajectories across the first 2 axes of this execution movement subspace (Figure 8). PG activity prior to the Go Cue was similar and overlapping for the two conditions and showed little variance in the movement subspace, reflected by the minimal evolution of the trajectories until this point. After the Go cue in execution, activity in each population then progressively evolved through different stages of the trial through HPR and DO, as indicated by the arrows, spanning the movement subspace (Figure 8A). During action observation, M1-PTNs (Figure 8A, top) and M1-UIDs (Figure 8A, middle) showed a highly collapsed trajectory, suggesting little similarity between population activity in execution and observation. F5 population activity, on the other hand, followed a qualitatively similar, but smaller trajectory to that seen during execution, with ordered progression through stages of the task (Figure 8A, bottom). For each population, we quantified the level of variance captured on these axes for both execution and observation. While the PCA method ensured that three dimensions captured the majority of the variance (>90%) of the execution data for all 3 populations (Figure 8B), captured observation variance was relatively low for both objects (5-10% for M1 populations, 20-25% for F5 for both objects). The ratio of this variance, to the maximum possible variance which could be captured within the observation data constituted a measure of alignment (Figure 8C, purple lines, see Methods). To quantify the significance of this overlap relative to what could be expected simply by chance, we compared this alignment to a null distribution of alignment calculated from pairs of random orthonormal dimensions. During movement, we found that only F5 showed an alignment between observation and execution greater than expected from chance for both grasps (PG alignment: 0.20, p = 0.003, WHG: 0.21, p < 1 × 10^−5^, upper-tailed permutation test). In M1-PTNs and M1-UIDs, on the other hand, alignment was not significantly different to chance (both grasps p > 0.1).

**Figure 8.**
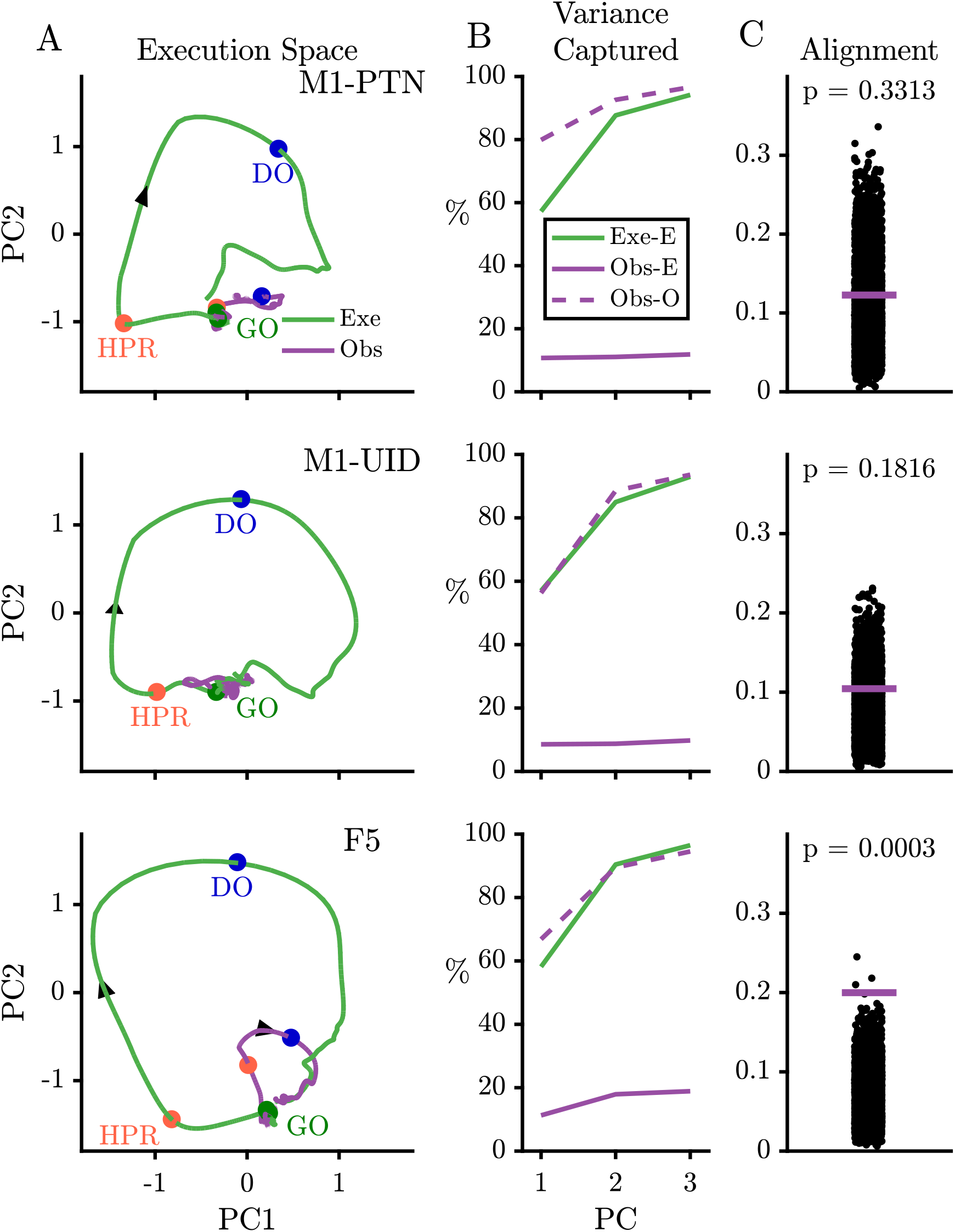
Execution and observation activity within a movement subspace. **(A).** Traces showing the evolution of M1-PTNs, M1-UIDs and F5 population activity within a 2-D movement subspace (defined by movement execution activity) across the whole trial during PG execution (green) and observation (purple) conditions. Larger coloured circles on each trajectory mark key events (Go, HPR, DO, labelled) in trial sequence which were used for multiple alignment of neural activity, and arrows on trajectories indicate direction of time. **(B).** Cumulative variance captured by the first three principal axes, separately for each sub-population. Exe-E (green), execution variance in execution subspace; Obs-E (purple), observation variance in observation subspace; Obs-O (dashed purple), observation variance in observation subspace. **(C).** Alignment index of observation activity in the movement subspace, for each population separately. Coloured horizontal line denotes observed alignment indices. Execution alignment index is equal to 1 by definition (not shown). Black points show alignment values from the null distribution.

### Movement suppression during action observation

Given that the patterns of neural activity show a clear divergence after the Go cue in the two conditions, we considered whether the activity during action observation contained signatures of movement suppression. We therefore examined the activity of the same populations of neurons during cued action withholding, and compared this to the responses during action observation. Figure 9A shows four single neurons recorded during PG execution, observation, and NoGo conditions. The activity of the M1-PTN and M1-UID (top panels) was clearly different for movement and non-movement around 100-150ms after the Go/NoGo cue, but showed comparatively little difference between observation and NoGo. By contrast, the activity of the M1-PTN in the lower left panel, which is the same neuron as shown in Figure 4A, was clearly different for all three conditions. The F5 neuron (Figure 9A, lower right) discharged in a similar way for execution and observation, first decreasing then increasing activity, while increasing activity in the NoGo condition. Using all neurons with at least 10 trials recorded per task condition, we trained a maximum correlation coefficient classifier to decode condition for each cortical population (Figure 9B). Across all three populations, the decoder was able to distinguish condition with high accuracy from 100-150ms after the Go/NoGo cue was given. We hypothesised that this could be largely driven by very reliable decoding of execution, which often shows greater variation in firing rates, and therefore also trained and tested the decoder with observation and NoGo conditions only (Figure 9B). This revealed a slower rise in accuracy, which was also different between the three populations. F5 showed significant decoding between these two conditions 300ms after the imperative cue, whereas for M1-UIDs and M1-PTNs, this was delayed until 400 and 450ms, respectively. Similar results were observed for decoding of WHG (in the 3-condition case, significant decoding occurred after 100-150ms, whereas when decoding between Observation and NoGo, significant decoding was again delayed; F5 – 300ms, M1-UIDs – 400ms, M1-PTNs 550ms). We also trained and tested the decoder on the other condition pairs (Execution-Observation, Observation-NoGo), and these also always produced strong decoding from 100-150ms after the Go/NoGo cue. Lastly, we performed a second PCA (Figure 10) this time defining each population’s subspace using observation activity after the Go cue (see Methods). We then projected each condition’s activity onto this subspace, which allowed us to compare the overlap of the execution and NoGo conditions with the observation subspace separately, in an analogous way to the analysis presented in Figure 8. In M1-PTN and M1-UID populations (Figure 10, first two rows), NoGo trajectories (orange) show a closer similarity to observation ones (purple). Although the M1-PTN population trajectory during NoGo condition showed smaller variance, its evolution over time was similar to the observation population trajectory, with the “trough” of both trajectories occurring at a similar time in advance of the average time of experimenter HPR (orange circles). By contrast, execution activity (green) showed quite different patterns to observation. In F5 (bottom row), the execution and NoGo trajectories both showed little variance, suggesting that neither condition overlap strongly with the observation subspace. Quantitatively (Figure 10C), M1-PTN NoGo activity overlapped with observation activity during this period significantly more often than chance (p = 0 for both grasps), and the raw alignment value was much larger for NoGo than for execution (PG: 0.31 vs. 0.06, WHG: 0.44 vs. 0.18). M1-UID NoGo activity also overlapped significantly with observation relative to the chance (both p = 0), whereas execution activity did not (both p > 0.1). By contrast, F5 NoGo and execution activity showed low levels of overlap with observation during this period, although this was significant relative to chance for WHG (0.03 and 0.03 for PG, both p > 0.3; 0.11 and 0.09 for WHG, both p < 0.0001).

**Figure 9.**
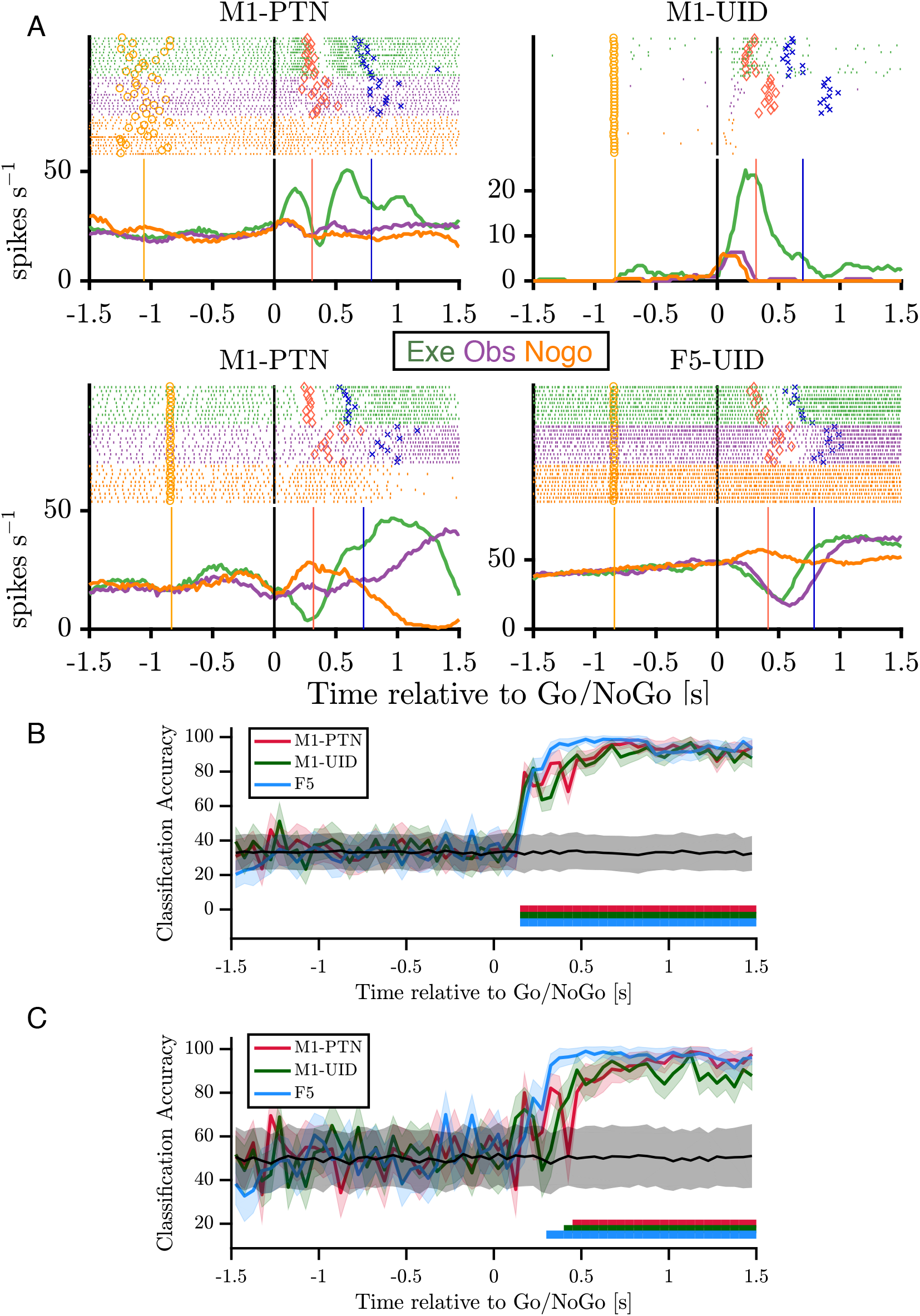
Activity during NoGo. **(A).** Example single-neuron responses during execution, observation, and NoGo. Each subplot shows a raster and histogram representation of single-neuron activity during PG execution (green), observation (purple), and NoGo (orange), with single alignment to the Go/NoGo cue (vertical black lines). Rasters and histograms are compiled from a randomly selected subset of 10 trials in each condition. For histograms, firing rates were computed in 20ms bins and boxcar-smoothed with a 200ms moving average. Event markers (LCDon, Object Cue etc.) are as shown previously (Figure 4). **(B).** Classification accuracy of maximum correlation coefficient classifier decoding between PG execution, observation, and NoGo conditions within each population. Grey trace and shading shows mean±1SD of decoding accuracy following permutation shuffling, and coloured bars along bottom show period of consistent significant decoding for each population. **(C).** As for (B) but decoding between observation and NoGo only.

**Figure 10.**
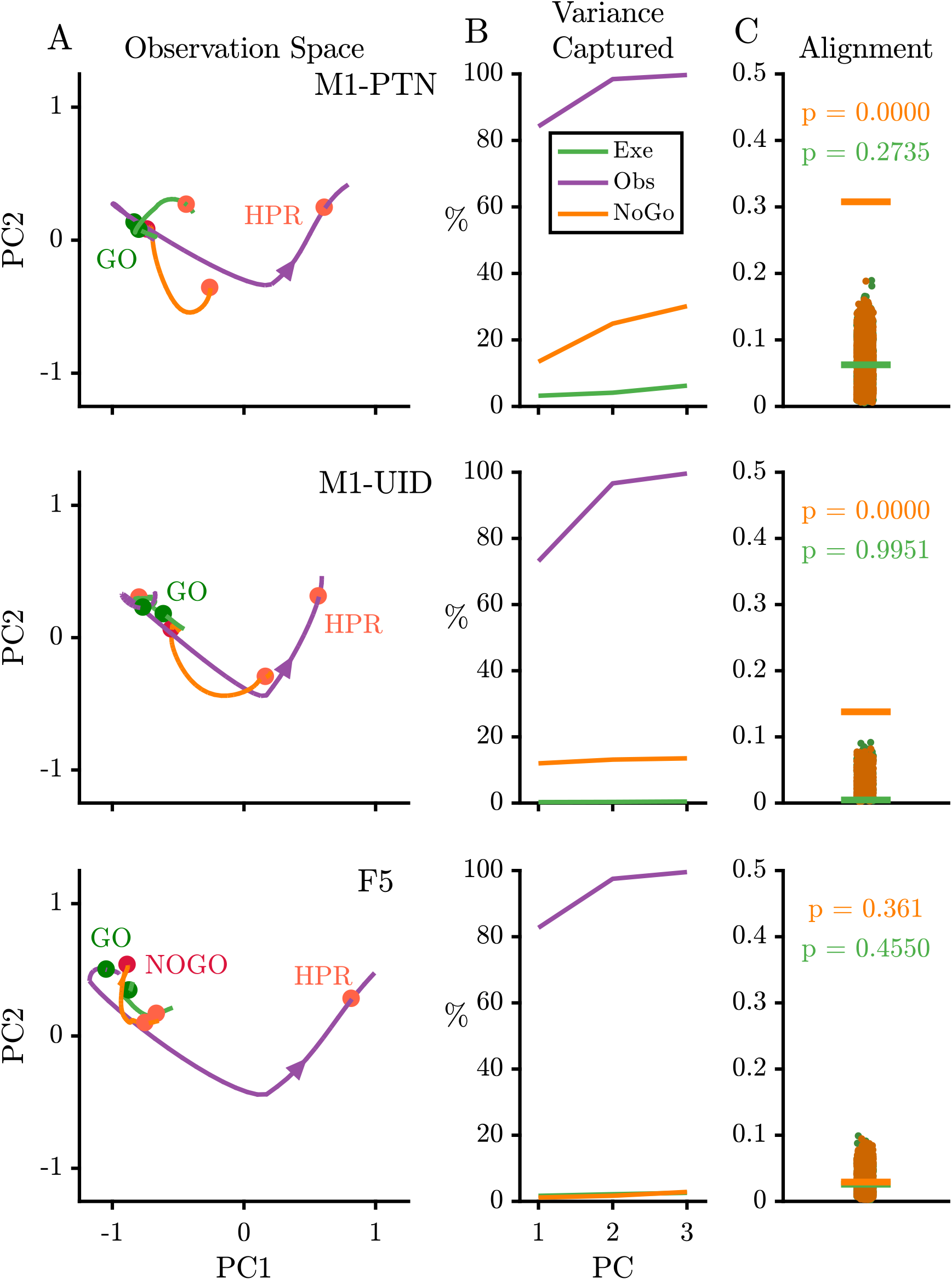
NoGo activity within an observation subspace. **(A).** Traces showing the evolution of M1-PTNs, M1-UIDs and F5 population activity during PG execution (green), observation (purple) and NoGo (orange) conditions within the first 2 dimensions of an observation subspace spanning the 100-400ms after the Go cue. Each trajectory show the −100 to +400ms period around the Go/NoGo cue (green/red circles). Average HPR time (across execution and observation) is also shown on each trajectory An arrow on observation trajectories indicates the direction of time. **(B).** Cumulative variance captured by the first three principal axes for execution, observation, and NoGo, separately for each sub-population. **(C).** Alignment indices of execution and NoGo activity in the observation subspace shown as coloured lines, for each population separately (Execution – green, NoGo – orange). The alignment index for observation activity is equal to 1 by definition (not shown). Scattered points show alignment values from the null distributions for execution and NoGo separately.

## Discussion

In this study, we investigated the relative contributions of F5 and M1 populations during the execution, observation, and withholding of grasping actions. We found that the modulation depth and profile of activity in F5 MirNs was more similar between execution and observation. In M1 populations, particularly M1-PTNs, although many neurons did modulate during both execution and observation, the magnitude and pattern of activity was distinct in these conditions, and observation activity shared parallels with activity when the monkeys simply withheld their own movement.

Previous interpretation of mirror activity has mostly been made in the context of known motor properties of the areas and pathways in question. F5 is critical for goal-directed visual guidance of the hand (Godschalk et al., 1981; Weinrich & Wise, 1982; Rizzolatti et al., 1998; Fogassi et al., 2001), and contains a ‘vocabulary’ of motor acts (Rizzolatti et al., 1988), supporting internal representation of different grasps (Murata et al., 1997; Raos et al., 2006; Umiltá et al., 2007; Spinks et al., 2008; Fluet et al., 2010; Schaffelhofer & Scherberger, 2016). F5 makes only a limited contribution to the corticospinal tract (CST) (Dum & Strick, 1991; He et al., 1993), but is anatomically (Muakkassa & Strick, 1979; Godschalk et al., 1984; Matelli et al., 1986; Dum & Strick, 2005), and functionally (Cerri et al., 2003; Shimazu et al., 2004; Schmidlin et al., 2008; Kraskov et al., 2011) strongly interconnected with M1. M1 provides the major drive to the CST and exerts a direct influence over distal hand musculature, which is probably exploited by executive commands necessary for control of skilled hand movements (Kakei et al., 1999; Brochier et al., 2004; Lemon, 2008).

### M1 observation activity is dissimilar to execution activity

We first confirmed that, although both F5 and M1 neurons can show mirror responses (Figure 4), F5 mirror activity during observation is more comparable in amplitude to execution activity (Figure 5). This is in line with previous reports of F5 MirN activity, suggesting a similar representation of grasp irrespective of whether the action is executed or observed (Gallese et al., 1996; Kraskov et al., 2009; Bonini et al., 2010). By contrast, M1 was first thought to completely lack MirNs (Gallese et al., 1996; Nelissen et al., 2005), and although several studies have now shown that neurons in this area, including PTNs, can show mirror responses, this activity is often relatively weak (Dushanova & Donoghue, 2010; Vigneswaran et al., 2013). Here, we found that M1-PTNs which increased firing during both execution and observation i.e. classical MirNs, showed a 3-4 times reduction in activity during observation relative to execution (Figure 5D), quantitatively comparable to previous reports (Dushanova & Donoghue, 2010; Vigneswaran et al., 2013). When examining the correlation of population-level activity across execution and observation, we found differences across different task periods, with M1-PTN MirNs showing a particularly weak correlation between the two conditions during the early stage of movement (Figure 6). Low-dimensional subspaces capturing variance associated with movement execution also captured meaningful observation variance in F5, but not in M1-UID and M1-PTN populations (Figure 8). It should be noted that the alignment of uniformly random orthonormal subspaces is dependent on the dimensionality, rather than structure, of the data, and therefore constitutes a relatively low bar for significance testing. However, an alternative method which seeks to circumvent this by constraining random subspaces to the covariance structure of the full dataset (Elsayed et al., 2016) is biased towards identifying orthogonality between two different subspaces, due to regression to the mean within the null distribution. The finding that execution and observation are more overlapping in F5 is consistent with recent work demonstrating MirN activity in ventral premotor cortex (PMv) and M1 during execution of reach and grasp to be associated with a series of hidden states, which were recapitulated during observation in PMv, but not M1 (Mazurek et al., 2018). Overall, although M1 neurons can be active during both execution and observation, the pattern of this activity at the population level was quite different between the two conditions. In a classical gating model of corticospinal control where increased activity in excitatory pyramidal cells drives movement, the net disfacilitation of M1-PTNs during observation provides a plausible substrate for inhibiting movement, given their anatomical and functional proximity to the spinal output (Kraskov et al., 2009; Vigneswaran et al., 2013). However, suppression of PTN activity has also been reported during movement execution tasks (Kraskov et al., 2009; Quallo et al., 2012; Vigneswaran et al., 2013; Soteropoulos, 2018), and was observed in the present task (Figure 4D). Suppression during movement could drive downstream inhibitory spinal circuits, given that PTNs not only make direct connections with motoneurons via the cortico-motoneuronal (CM) system (Lemon, 2008; Rathelot & Strick, 2009), but also connect to segmental interneurons within the spinal cord (Kuypers, 1981), and tightly timed suppression of muscle activity is essential for skilled movement (Quallo et al., 2012). An alternative, but not mutually exclusive, possibility, is that population activity at the cortical level evolves within a dynamical system, which implicitly gates downstream circuitry (Kaufman et al., 2013, 2014; Elsayed et al., 2016), although this framework is not yet reconciled with the known anatomy of neuronal sub-populations. Under the assumption that the balance of excitation and inhibition at the motor cortical level is fundamental for movement generation and suppression, then it should be expected that the respective patterns of activity during execution and observation should be reflected in the resultant behaviour. From both a representational and dynamical systems viewpoint, M1 activity during execution and observation, particularly in PTNs, may be sufficiently dissimilar so as to ensure movement is only produced in the former condition.

### Correlates of movement suppression in M1 observation activity

The dissociation between execution and observation appeared most prominent around the time of movement onset, in line with previous suggestions regarding the role of MirNs in movement suppression (Kraskov et al., 2009; Vigneswaran et al., 2013). In the present study, we used a pseudo-randomised trial sequence, and Go/NoGo and execution/observation information was provided simultaneously on each trial (Go/NoGo cue; Figure 1B). This contrasts with most action observation studies in which block-designs are used, and may provide a more ethological framework for assessing functions of the CST in movement suppression. We identified movement-related cortical neurons responding to both observation and NoGo conditions to varying degrees (Figure 9A). A decoder trained to discriminate between three conditions exceeded chance and reached plateau 100-150ms after the Go/NoGo cue (Figure 9B), presumably the time necessary for visual information about trial type to become available to motor areas. A second decoder trained to distinguish only between observation and NoGo took longer to exceed chance performance, particularly for M1 neurons, indicative of similar activity patterns in the two conditions (Figure 9C). This was corroborated by analysis of the evolution of activity within an observation subspace after the Go cue, which captured significant NoGo variance in M1-PTNs, but less so in F5 (Figure 10). Taken together, these results suggest a greater overlap between observation and NoGo neural states in M1 than F5. The task design itself, with observed actions taking place within the monkey’s reach, may introduce some similarity between observation and NoGo, such that the strategy adopted by the monkeys is to treat the observation Go cue and NoGo cue similarly. Observed actions occurring in peri-personal space often modulate MirN responses differently to when the action is beyond the monkey’s reach (Caggiano et al., 2009; Bonini et al., 2014a; Maranesi et al., 2017), suggesting the capability to interact with observed actions is an important aspect of mirror activity. A further important point is the difference between F5 and M1, which indicate that while M1’s priority is to distinguish movement from non-movement from an egocentric perspective, F5 maintains a more similar representation across executed and observed actions, independent of the acting agent’s identity. These results suggest the formulation of a simple model framework, in which the movement execution and suppression-like features of the unfolding action observation response in M1 (and F5) reflect a balance of the activity patterns seen during the execution and NoGo conditions. This balance could be determined by inputs from upstream areas within the MirN system, and prefrontal areas responsible for encoding general features of action and self versus other encoding, as well as intrinsic dynamics within premotor and motor cortex. State-space analyses provide a useful tool for analysing these temporal dynamics during different stages of action execution, observation, and withholding. Future investigations which more widely sample grasping execution state space (i.e. recording from more neurons but also using a much more extensive range of movement and grasping conditions) may be able to address this, and also relate neural correlates of suppression during observation and NoGo conditions to the withholding of movement during instructed delay periods (Kaufman et al., 2014; Elsayed et al., 2016; Hannah et al., 2018; Soteropoulos, 2018). Single-trial analyses may also hold particular relevance for examining the switching of neural state between initiation and suppression of movement in the context of an action execution-observation task. Our dataset, with small samples of recorded cells per session, was not well suited to this type of analysis.

The timing and kinematics of monkey and experimenter movements were clearly different, which could explain why similarity between execution and observation decreased during the reaching phase, however, there are several reasons this is unlikely to be a dominant factor. Firstly, correlations between execution and observation already began to decrease during the late reaction period, i.e. before any movement had occurred (Figure 6B). At the single-neuron level, firing rates showed little correlation with movement speed (inversely proportional to movement time given constant distance between hand and objects) (see also Vigneswaran et al., 2013). Furthermore, given that many sessions involved simultaneous recording of units in F5 and M1, timing reasons could not explain differences between the sub-populations. The targeting of recordings to F5, an area with a preponderance of grasp-related activity (Rizzolatti et al., 1988; Gallese et al., 1996; Raos et al., 2006; Umiltá et al., 2007; Michaels et al., 2018), and the M1 hand area, may also contribute to closer similarity between execution and observation during grasp and hold, rather than reach periods of the task. However, we did not impose strong online selection criteria regarding the proximal vs. distal related activity of recorded cells (in particular, all stable and well-isolated PTNs, once identified, were recorded for a full set of trials), and rICMS at some recording sites elicited movements of proximal muscles. This is also consistent with a developing body of literature involving anatomical tracing, stimulation mapping and task-related activity which questions the simple segregation of dorsal and ventral premotor cortex into reaching and grasping areas, respectively (Raos et al., 2003, 2004; Dum & Strick, 2005; Stark et al., 2007; Lehmann & Scherberger, 2013; Takahashi et al., 2017). On the other hand, although there is now ample evidence that cells in dorsal premotor areas, or within proximal limb representations in M1, do mirror reaching movements (Cisek & Kalaska, 2004; Dushanova & Donoghue, 2010; Papadourakis & Raos, 2019), to our knowledge, the anatomical identity of these MirNs, and their potential influence on downstream targets, has not been directly tested, and could be of particular relevance for initiation or suppression of reaching movements.

Overall, the results of this study point to a functional distinction in premotor and motor cortex respectively regarding the representation of executed and observed grasping actions. F5 neurons appear more engaged in the encoding of the upcoming grasping action, such that execution and observation activity remain similar over a longer time course. Contrastingly, M1 populations, and M1-PTNs in particular, show a more flexible dissociation through the task, first distinguishing movement from non-movement, which may then allow subsequent grasp-related mirror activity to evolve in the absence of self-movement.

## Conclusions

In this study, we confirm that F5 activity is closer in amplitude and profile during action execution and observation, whereas M1 showed a particularly weak relationship in activity between the two conditions. The M1 neural state during observation diverges from the execution state in the lead-up to movement onset, and contains signatures of an action withholding state at this time. Functionally, the different patterns of activity between execution and observation in the two areas could support a context-dependent dissociation between grasp-related visuomotor transformations and the recruitment of descending pathways for elaboration into actual performance of skilled grasp. The increasing capabilities for wide-scale simultaneous recordings from many neurons, and accompanying inactivation and manipulation experiments, should help to shed further light on the transfer of information through premotor and motor areas for the representation and organisation of goal-directed actions and the observation of these actions.

## Acknowledgements

The authors thank Tabatha Lawton, Dominika Klisko, and Adam Keeler for help with recordings, and Spencer Neal, Jonathon Henton, Chris Seers, and Martin Lawton for technical assistance. Roger Lemon provided useful feedback on an earlier version of the manuscript.

## Notes

**Conflicts of Interest:** The authors declare no competing financial interests.

**Funding:** S.J.J was funded by a Brain Research UK Graduate Student Fellowship. A.K. was funded by the Wellcome Trust, grant number 102849/Z/13/Z.

